# SPINK1-induced amelioration of impaired autophagy contributes to suppression of trypsinogen activation in a model of acute pancreatitis

**DOI:** 10.1101/2020.03.06.981654

**Authors:** Yuxiao Zhao, Jianlong Jia, Abdullah Shopit, Yang Liu, Jun Wang

## Abstract

SPINK1 has been regarded as a reversible trypsinogen inhibitor for the inappropriate activation of trypsin, a key step in the initiation of acute pancreatitis (AP). However, the mechanisms of its action remains largely unclear and controversial. Here, we reported an unexpected effects of SPINK1 on inhibiting trypsinogen activation through the regulation of impaired autophagy in cerulein-stimulated AR42J cells, a well-established in vitro model of acute pancreatitis. Firstly, we found that the impaired autophagic flux was induced and trypsinogen activity enhanced in the above setting. Then, we showed that SPINK1 overexpression could inhibit the level of increased autophagic activity, improving the hindered autophagy flux, and significantly decreased the trypsinogen activity, whereas shRNA-caused downregulation of SPINK1 exacerbated the impairment of autophagic flux and trypsin activity, in the same cerulein-processed cells. More importantly, the trypsinogen activation in this model could be ameliorated by 3-Methyladenine(3-MA), an autophagy inhibitor. Thus, this study revealed, possibly for the first time, that SPINK1 greatly blocked the trypsinogen activation possibly through the modulation of impaired autophagy in cerulein-induced in vitro model of acute pancreatitis.

## 1. Introduction

The mechanisms in the pathogenesis of acute pancreatitis (AP) remains incompletely understood. SPINK1, serine protease inhibitor Kazal-type 1, was originally identified as a reversible inhibitor of accumulated active trypsin produced in the triggering step of AP [1]. Although SPINK1 was found to be responsible for the negative regulation of autophagy in normal pancreatic acinar cells [2], it’s roles spanning over autophagy and trypsinogen activation are still kept undiscovered in the development of AP. Within the past decade, the impaired autophagic activity is thought to be possibly involved in the initial episode of acute pancreatitis, though the underlying principles are elusive[3]. The appearance of vacuoles inside acinar cells features an early stage of development in different models of acute pancreatitis, and these vacuoles are considered to be autophagic in nature[4, 5]. Interestingly, the numerous zymogen granules are found to be accumulated within these autophagic vacuoles, autophagosome or autophagolysosomes, on the onset and early course of experimental AP[2, 5], suggesting the possible connection between autophagy and proteinase activation. Besides, the amount of long-lived protein, an indicator of disordered autophagy flux, was increased in AP setting [6]. Of note, the loss of SPINK1 in mice could give rise to enhanced autophagy in the acinar cells of normal pancreas [2], and the targeting overexpression of rat PSTI-1(SPINK1) to the pancreas in transgenic mice significantly reduced the histologic severity of AP, and the trypsin activity was also significantly lower in the above mice receiving cerulein injections[7]. Thus, the occurrence of autophagy in AP was likely associated with activation of the zymogen granules, while SPINK1’s presence was possibly related to regulation of the autophagic event of the disease[8]. In acute pancreatitis, it is generally accepted that the premature and intrapancreatic activation of trypsinogen is an early triggering step for the disease[1, 4]. However, the mechanisms or intermediate links by which SPINK1 blocks trypsin activation in the pancreas remains largely unknown. Based on the aforementioned data, therefore, we assumed that SPINK1 may inhibit the active trypsin likely through the regulation of autophagy in the initiation of AP of a cellular model.

## 2. Materials and methods

### 2.1. Cell culture, reagents and vectors

AR42J cells, indirectly purchased from ATCC, were cultured in F12K medium supplemented with 10% FBS. Cerulein is obtained from MedChem Express, NJ; (CBZ-Ile-Pro-Arg)_2_-R110 (R6505), a substrate for trypsin, from Molecular Probes, Inc., Invitrogen, OR; 3-Methyladenine (3-MA), from MedChemExpress, NJ; Boc-Gln-Ala-Arg-AMC, substrate for trypsin, from Enzo Life, Inc. NY. All other chemicals of highest purity were commercially available and obtained from either local or domestic Chemicals Ltd. Rat SPINK1 monoclonal antibody (D264388) and anti-MAP1LC3B polyclonal antibody(D163557) were purchased from Sangon Biotech, Shanghai, China. pLenti-CMV-hSPINK1-2A-GFP vector(rat) was obtained from Vigene Biosciences, Shandong; pilenti-SPINK1-shRNA-GFP-Puro construct(rat) from Applied Biological Materials(abm), Nanjing.

### 2.2. Cerulein-induced in vitro acute pancreatitis

Briefly, AR42J cells were seeded into culture plates 24 hours before they were stimulated with cerulein (10 nM) for different time periods, respectively[9].

### 2.3. Measurement of intra-acinar trypsin activity in vitro

For living cells, the intra-acinar trypsin activity was measured in 1× 10^4^ overnight-culturing AR42J cells/per well that were stimulated with 10 nM cerulein for varied time duration in 96-well plate. The analysis protocol was modified based on the instruction sheet for Rhodamine 110-based proteinase substrates from Molecular Probes, Invitrogen. 10 µM Rhodamine 110 – based trypsin substrates (CBZ-Ile-Pro-Arg)_2_-R110(R6505) was added into the cells for 15 min at 37°C before performing the measurement. The multifunctional fluorescence reader (BioTek Synergy NEO, Germany) was set with peak excitation and emission wavelengths of 498 nm and 521 nm, respectively. For lysed cells, the measurement protocol for trypsin activity was the same as described above except that the culturing medium in the well was replaced by cell lysis buffer, and then the cells were intermittently stirred around 20 min by a mixer for complete dissolvement before carrying out the measurement. The components of lysis buffer are: 50 mmol/L Tris-HCL, pH 8.0; 150 mmol/L NaCl; 1 mmol/L EDTA; 0.1% SDS. All the experiments were carried out in triplicates for each treatment and carried out at lease two times.

### 2.4. Transfection and transduction of AR42J cells

The AR42J cells were transfected or transduced with pCAG-pHluorin-TagRFP-mLC3B plasmids and SPINK1-GFP and SPINK1-shRNA lentiviral particles, and the scrambled or respective vectors were used as a negative control. The transfection or transduction procedures were carried out accordingly, based on the instruction manual. The transfection or transduction efficiency was 40%∼60%, and assessed by the ratio of positive GFP cells over the total, via the observation under the fluorescence microscope 48 hrs after transfection or transduction.

### 2.5. Measurement of proliferation rate and cellular colony formation

Cell proliferation rate was determined using Cell Counting Kit-8 (CCK-8). Briefly, AR42J cell’s suspension treated with or without cerulein (10 nM) or/and 3-MA (10 mM) was inoculated in a 96-well plate (0.5 × 10^4^/well). 10 μl CCK-8 solution were added into each well and the incubation was continued for additional 2-4 hrs. The experiment was performed in triplicates for each group using a 96-well plate. The absorbance (A) in the wells was measured using a micro-plate reader (iMark, BIO RAD, Japan). The 2500 SPINK1-overexpressing cells of single cellular suspension were plated in 6-well plates and cultured for **7** days, and then these cells were stained with 0.005% (w/v) crystal violet in 25% (v/v) methanol before they were counted under a microscope. The proliferation rates were calculated as: growth rate (%) = (1 – treated group A/control A) × 100%.

### 2.6. Evaluation of autophagic flux

At first, the autophagic flux in in vitro acute pancreatitis was evaluated by the expression patterns of LC3II, a typical autophagy marker and P62, a substrate of autophagy[10], in the respective cerulein-stimulated AR42J cells using Western blotting. Secondly, the transfection of dual-Fluorescence pCAG-pHluorin-TagRFP-mLC3B construct[11] into the above AR42J cells was used for the examination of the ratio of yellow/red fluorescence puncta via the method described below. Briefly, the correspondingly transfected AR42J cells were incubated on the coverslips that were placed on the bottoms of a 6-well plate with complete media for 48 hours before they were observed under fluorescence microscope (IX 71, Olympus, Japan). Oil immersion lens was applied for the observation of green and red bright puncta in SPINK1 overexpressing or knockdown AR42J cells stimulated with cerulein. The number of bright fluorescence spots within about 80 cells of 10 fields were counted three times under the fluorescence microscope. The merged yellow spots from the green and the red indicated the presence of autophagosome while the red ones only standed for the autophagolysosome since the green fluorescence quenched within the lysosome at low pH. For the estimation of autophagy flux state, the average ratio of yellow over red fluorescence puncta was calculated from two independent experiments.

### 2.7. Western blotting

Western blot analysis was carried out according to the method described previously [13]. Primary antibodies were diluted at indicated concentration for the following antigens:SPINK1, 1:1000; LC3B, 1:1000; ATG5, 1:1000; p62, 1:1000. The secondary antibodies(Sangon Biotech, Shanghai) to detect all proteins were used at varied dilution ratio.

### 2.8. Statistical analysis

The quantitative or semi-quantitative variables were counted as mean ± SEM and the means were compared correspondingly using unpaired two-tailed Student’s *t* test. The statistical significance was assigned to *P* values < 0.05.

## 3. Results

### 3.1. Accumulation of intra-acinar active trypsin in in vitro cerulein-induced acute pancreatitis

To assess whether in vitro model of AP were successfully generated, the intra-acinar trypsinogen activity were examined in cerulein-stimulated AR42J cells originally derived from rat tumorigenic exocrine pancreas. Compared with controls, the intra-acinar trypsinogen activity in the alive cells, a key trigger in the initiation of AP, began to significantly elevate at 5 min after cerulein stimulation (10 nM), and that value gradually raised up to the maximum at 30 min in living AR42J cells, followed by a gradually decreased levels within the next 5-6 hrs (Fig. 1A, P < 0.05 ∼ 0.001). For sureness, we also measured the trypsinogen activity in the lysates of cerulein-treated AR42J cells, showing the increased levels of active trypsin at 30, 60 min post stimulation as well (Fig. 1B). The above data suggests that a successful setup of in vitro model of AP in our study.

**Fig.1.**
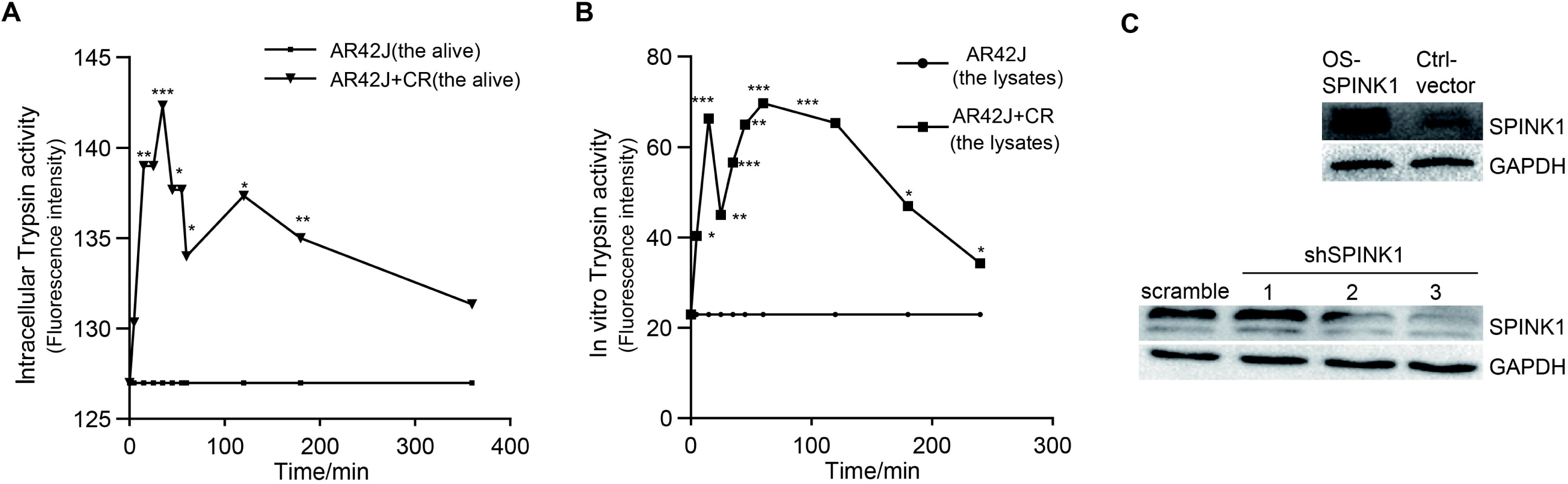
Accumulation of intra-acinar active trypsin in ex vivo acute pancreatitis and SPINK1-related transduction of AR42J cells. (A) Measurement of trypsin activity in living cells after cerulein (CR, 10 nM) treatment. (B) Measurement of trypsin activity in AR42J cells lysate after CR (10 nM) stimulation. (C) Western blotting for SPINK1 over-expression and SPNK1 knockdown (sh-SPINK1, sh-SPINK2, sh-SPINK3) in lentivirally transduced AR42 cell lines, compared to controls. (n = 3 independent samples, Data represents the mean ± S.D. of triplicate experiment; *P ≤ 0.05, **P ≤ 0.01, ***P ≤ 0.001).

### 3.2. Generation of SPINK1 overexpressing and knockdown AR42J cell lines

For the purpose of investigation, the lentivirally transduced stable AR42J cell lines overexpressing (OS) and silencing SPINK1 were generated based on the protocol provided by Applied Biological Materials Inc.(abm, BC, Canada). Briefly, pLenti-CMV-hSPINK1-2A-GFP construct or pLenti-SPINK1-shRNA-GFP-Puro **v**ector with the ‘highest’ (Fig. 1C) was co-transfected with the mixture of pMD2.G and psPAX2 /packaging plasmids (Addgene Inc, Cambridge, MA, USA) and Lipo2000 transfectant (Invitrogen Inc.) into 293T cells, while the transfected empty or scrambled vectors was used as a negative control. The strong GFP fluorescence could be observed under the fluorescence microscope 24-48 hours after the transfection. The lentiviral particles were collected at 24 and 48 hrs after virus packaging. The AR42J cells were infected by the filtered lentiviral particles, and those transduced cells were screened under the pressure of 2 µg/ml puromycin for about 3-4 weeks before the drug-resistant cells were collected and passaged. Then, the protein expression of SPINK1 was measured in the passaged AR42J cell lines using Western blotting, showing that SPINK1 expression was correctly overexpressed or silenced in the corresponding AR42J cell lines (Fig. 1C).

### 3.3. Impaired Autophagy flux in acute pancreatitis ex vivo

To evaluate the state of autophagy flux in cerulein-stimulated AR42J pancreatic acinar cells, we first examined the expression of LC3I-derived LC3II, a typical marker of autophagy, and Western blotting analysis showed that LC3II was significantly upregulated in cerulein-processed AR42J cells at 8 hrs post-stimulation (Fig. 2D). We also examined the expression of p62, a substrate of autophagy, in the above-mentioned scenario. In contrast, p62 levels were significantly elevated at 8 hrs post-cerulein stimulation in cerulein processed AR42J cells(Fig. 2D). In consideration of reverse changing pattern of LC3B and p62 expression, these data suggested that autophagy flux possibly was hindered in this AP setting, since LC3B expression was upregulated whereas p62 amounts was not correspondingly consumed less as a autophagic substrate[10]. To further approve this finding, we pre-transfected pCAG-pHluorin-TagRFP-mLC3B plasmid, a dual-fluorescence(Green and red) construct used for evaluation of autophagy flux, into cerulein-treated AR42J cells, and counted the number of autophagosomes (merged yellow fluorescence spots) and autophagolysosomes (red fluorescence puncta) observed under a fluorescence microscope (Fig. 2A) [11]. Compared with the group without cerulein, the ratio of the yellow/red spots significantly increased in the group upon cerulein stimulation alone, indicating the presence of impaired autophagy flux in in vitro AP model, as the less formation of autophagolysosomes (red spots) implied a blockage of lysosome fusion and degradation at the early stage of AP (Fig. 2B, C, P < 0.01 ∼ 0.001). In contrast, SPINK1 overexpression could reverse this trend by increasing the ratio of red/yellow spots and SPINK1 knockdown could decrease this ratio even further (Fig. 2B, C, P < 0.01), demonstrating that SPINK1 was capable of ameliorating the impaired autophagy flux in the in vitro model of AP. Collectively, SPINK1 modulated the dysregulated autophagy flux in the aforementioned disease setting.

**Fig. 2.**
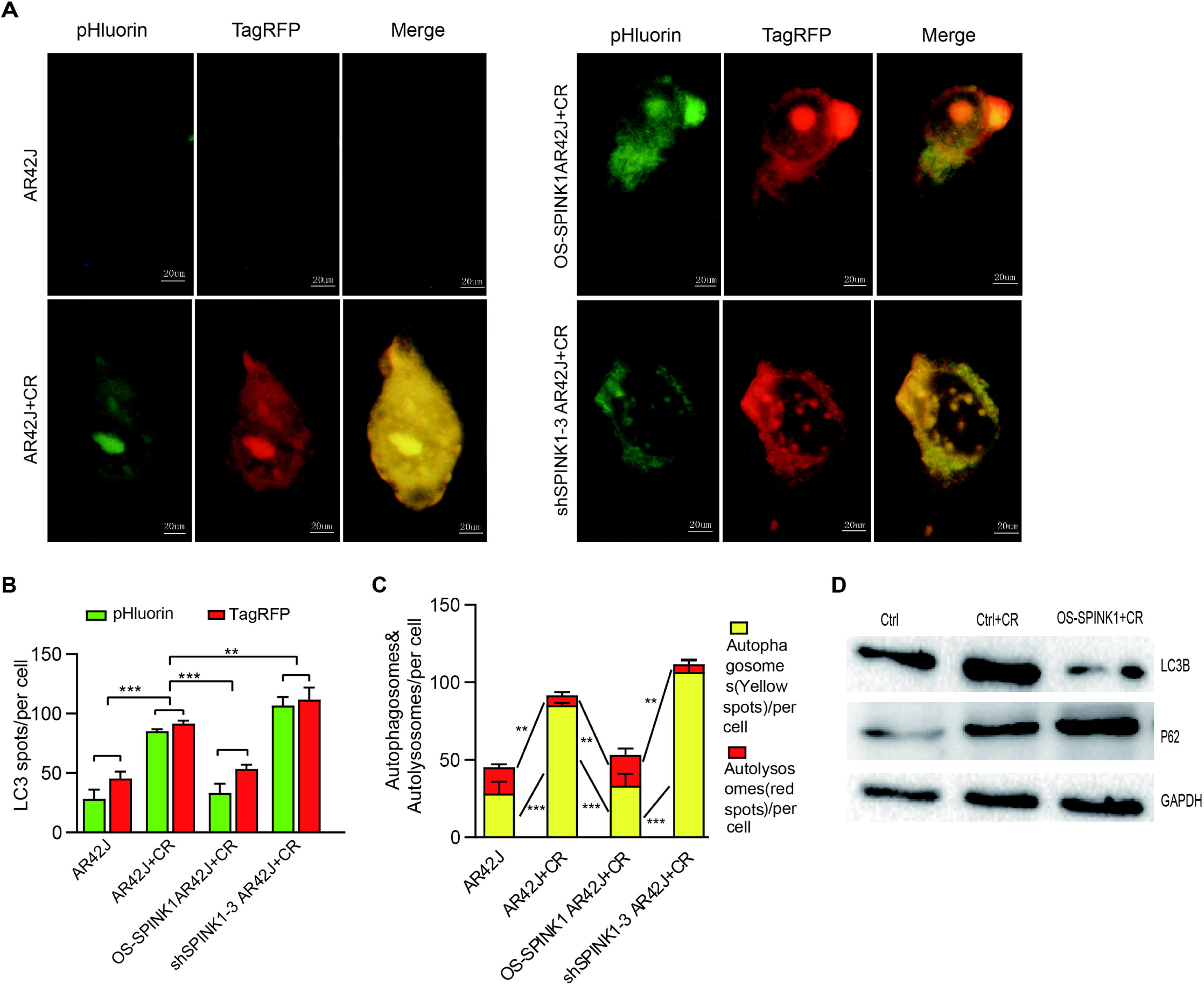
Effects of SPINK1 on autophagy flux in CR-induced acute pancreatitis ex vivo. (A) Observation of pCAG-pHluorin(the green) and TagRFP-mLC3B (the red) expressions under fluorescence microscopy, in empty, SPINK1-OS and SPINK1-knockdown cells, with or without CR treatment; Merger of green and red fluorescence spots as yellow puncta using the software. (B) Numbers of pHluorin relative to TagRFP-LC3 spots in corresponding cells. (C) Number of TagRFP-LC3 versus merged yellow spots in respective cells. (D) Expressions of LC3B and P62 in CR-treated cells overexpressing(OS) SPINK1. (Data represents mean ± S.D. of three independent times of calculation, *P ≤ 0.05, **P ≤ 0.01, ***P ≤ 0.001; scale bar, 20 uM).

### 3.4. Regulation of trypsinogen activity by autophagy state in in vitro acute pancreatitis

Since both altered autophagy state and trypsinogen activation were observed in in vitro model of experimental AP, we aimed to investigate the possible relationship between autophagy and trypsinogen activation in this setting. To approve this assumption, we used 3-MA(3-methyladenine), an inhibitor of PI3K and autophagosome formation, to suppress the autophagic event and found that the cerulein-induced trypsin activity was remarkably suppressed with the addition of 3-MA (10 nM) (Fig. 3A, P < 0.01). In short, the data suggested that the trypsin activity could be possibly modulated via the autophagic changes in experimental AP model.

**Fig. 3.**
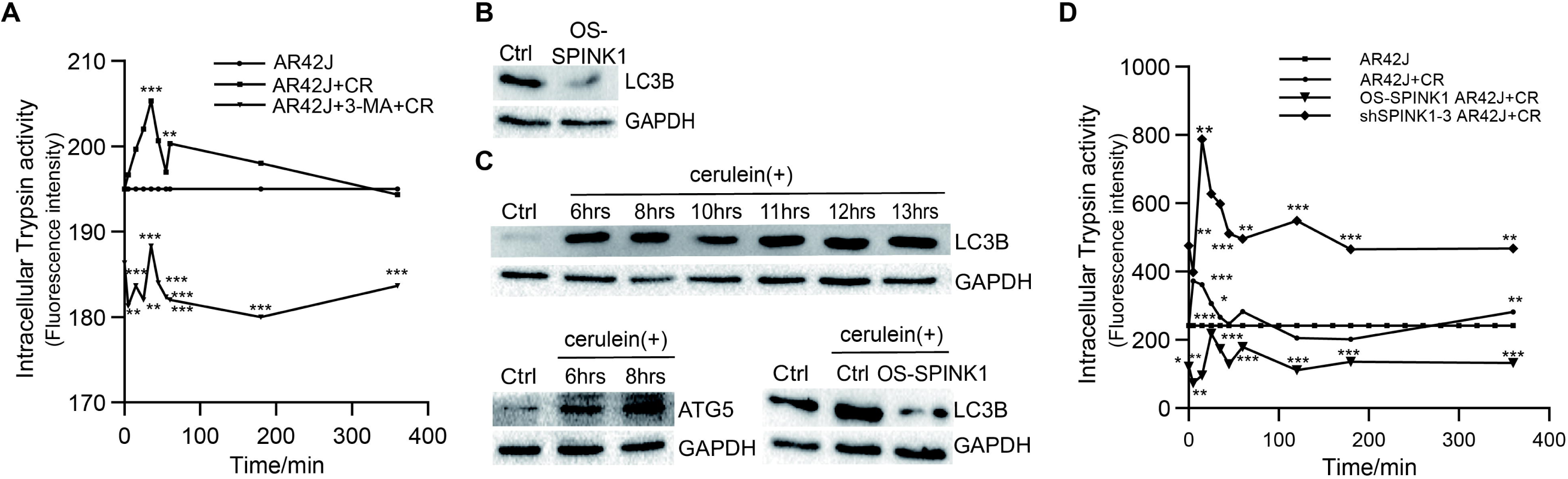
Expression of autophagic proteins, autophagy and SPINK1 regulation of trypsin activity in CR-induced in vitro acute pancreatitis. (A) Effect of 3-MA on intracellular trypsin activity of CR-stimulated AR42J cells. (B) Western blot analysis for LC3B in SPINK1 overexpressing AR42J cells without CR stimulation. (C) LC3B and ATG5 expressions in CR-treated cells at different times using Western blot. (D) Effect of SPINK1 on intracellular trypsin activity in CR-induced acute pancreatitis ex vivo. (n = 3 independent samples, data represent mean ± S.D. of triplicate experiment; *P ≤ 0.05, **P ≤ 0.01, ***P ≤ 0.001).

### 3.5. Decreased autophagic activity associated with overexpression of SPINK1

To study the effects of SPINK1 on autophagy, Western blotting was carried out in SPINK1 permanently overexpressing AR42J cell lines with or without cerulein stimulation. The expression of LC3B, a typical autophagy marker, was inhibited significantly in SPINK1 overexpression AR42J cells (Fig. 3B). To explore the effects of SPINK1 on the impaired autophagy induced by cerulein at the early stage of acute pancreatitis, we performed Western blotting and examined the changing patterns of ATG5 and LC3B in this scenario. Of note, compared to control, the expression of ATG5 and LC3B, two important biomarkers of autophagy, were significantly elevated in cerulein-stimulated AR42J cells at varied time points (Fig. 3C). However, the permanently SPINK1-overexpressing AR42J cells remarkably inhibited the expression of LC3B (Fig. 3C). Therefore, SPINK1 upregulation likely ameliorated autophagy activity in AR42J cells with or without cerulein treatment, i. e., both in normal growth or in in vitro state of AP.

### 3.6. SPINK1 regulation on accumulation of intra-acinar active trypsin in in vitro acute pancreatitis

To probe the relationship between SPINK1 and trypsin activity in the triggering step of AP setting, we examined the levels of active trypsin in SPINK1-OS/silencing AR42J cells upon cerulein-treatment at varied time points post-stimulation with cerulein. Relative to the increased trypsinogen activity upon cerulein administration, the ectopic SPINK1 overexpression significantly decreased trypsin activity and SPINK1 knockdown, in contrast, attenuated the extent of active trypsin, indicating that SPINK1 possessed the negative regulatory role in the maintenance of appropriate amount of active trypsin in the initiation of experimental acute pancreatitis (Fig. 3D, P < 0.01). Altogether, SPINK1 could regulate the degree of trypsinogen activity induced in cerulein-stimulated AP model.

### 3.7. Influence of SPINK1 on clonogenic survival and proliferation of pancreatic Acinar cells without cerulein treatment

To evaluate the effects of SPINK1 on biological phenotypes of AR42J cells, we performed CCK-8 and colony formation assays. Compared with empty-vector control, the metabolic and proliferative capability measured increased significantly in SPINK1 overexpressing AR42J cells, and decreased obviously in the above SPINK1 knockdown cells (Fig. 4C, P < 0.001). Noticeably, the clonogenic assay showed that the number of colonies formed in SPINK1-overexpressed AR42J cells was greater than those of empty-vector control, whereas SPINK1 knockdown cells remarkably fewer than control AR42J (Fig. 4A, B, P < 0.001). In summary, these data suggested that SPINK1 possessed the ability to promote the proliferation and clonogenesis of normal AR42J cells.

**Fig. 4.**
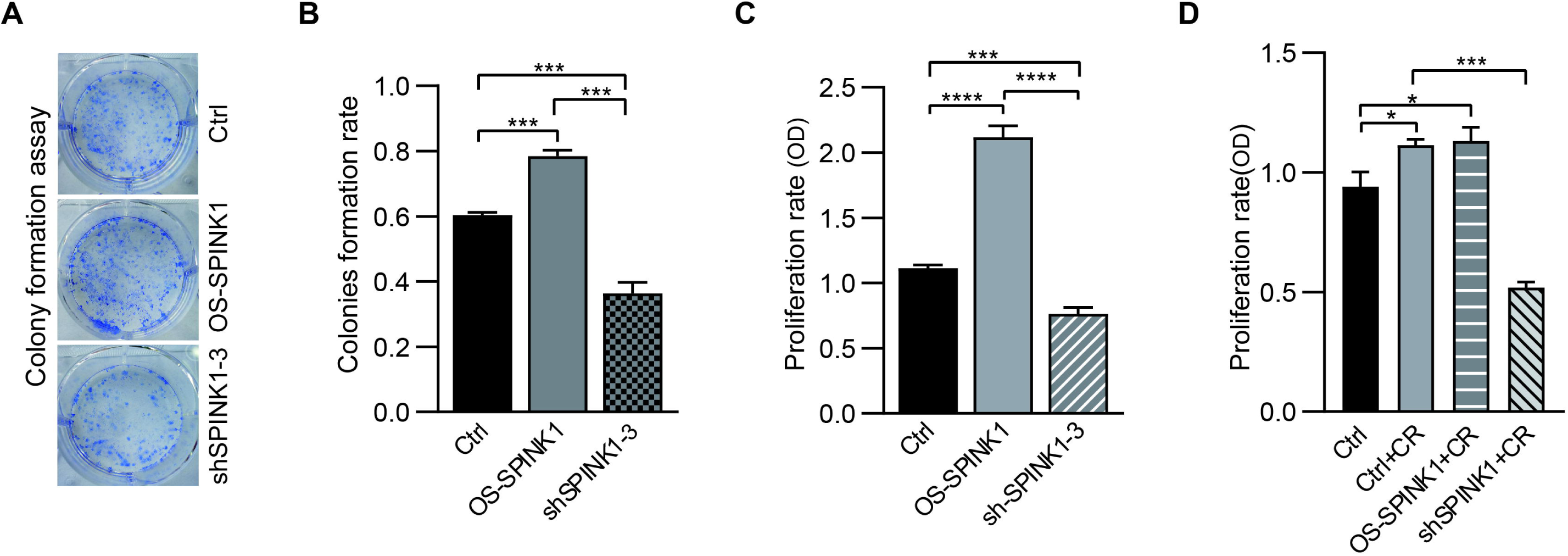
Influence of SPINK1 and cerulein on cellular proliferation and/or clone formation. (A) (B) Colony formation assay for SPINK1-overexpressing(OS) and knockdown AR42J cells. (C) Measurement of cell proliferation for CR-untreated SPINK1-OS and knockdown cells using CCK-8. (D) CCK-8 proliferation assay for CR-stimulated SPINK1-OS and silencing cells. (n=3 independent samples; data represent mean ± S.D. of triplicate experiment; *P ≤ 0.05, **P ≤ 0.01, ***P ≤ 0.001).

### 3.8. Influences of SPINK1 on proliferation of pancreatic AR42J acinar cells with cerulein stimulation

To explore the roles of SPINK1 in influencing biological features of AR42J cells in the circumstance of acute pancreatitis, we next measured the proliferative rates of the corresponding AR42J cell lines upon cerulein treatment. The result showed that the proliferation rate of cerulein-treated parental AR42J cells slightly increased at 8 hrs post stimulation (Fig. 4D, P < 0.05), though we are not aware of the mechanisms for the increment. Relative to the group without cerulein, the proliferative capacity began to upgrade in the cerulein-treated SPINK1 overexpressing AR42J cell line(Fig. 4D, P < 0.05), even greater than the treated parental AR42J (Fig. 4D, P > 0.05), in reverse to that decreased rate observed in SPINK1 knockdown AR42J cells (Fig. 4D, P < 0.001). These data suggested that SPINK1 could clearly exert some beneficial effects on the phenotypic state of pancreatic acinar cells of acute pancreatitis. This positive impact of SPINK1 might possibly help the pancreatic acinar cell fighting against the pathologic damage of pancreatic cells of acute pancreatitis.

## 4.0. Discussion

In this study, we demonstrated that SPINK1 could inhibit intra-acinar trypsin activation via amelioration of impaired autophagy in cellular model of in vitro acute pancreatitis, a well-accepted AR42J-related AP setting nowadays. Since the past decade, researchers have noticed that impaired autophagy was existed during the development of acute pancreatitis[14, 15], though the mechanisms was unraveled. On the other hand, SPINK1, as an active trypsin inhibitor, has been found to be a negative regulator of autophagy in normal pancreatic acinar cell, and the slightly enhanced trypsin activity can be detected in the isolated pancreatic acinar cells of SPINK1 knockout mouse pups [2, 16]. However, we are unaware of the underlined mutual connection among these events and the molecular mechanisms involved in particular, and as well as its significance in the development of the disease[17].

Here, firstly, our data demonstrated that the hindered autophagy flux was generated upon cerulein stimulation of pancreatic acinar AR42J cells, since it has been a controversial issue whether autophagy promotes or protects against the development of acute pancreatitis [18, 19, 20]. We showed several pieces of evidence that supported the above temporary conclusion. including elevated LC3B versus increased expression of p62, an autophagy substrate. These reversed trend of expression of LC3B versus P62 indicated that autophagy flux was partially damaged in experimental AP setting. For further approvement, the double fluorescence vector was applied to display the presence of lowered ratio of autophagosomes/autophagolysosomes (yellow/red spots) in cerulein-stimulated AR42J cells. The overexpression of SPINK1 could effectively upgraded the ratio of yellow/red fluorescence puncta, demonstrated the amelioration of SPINK1 on the impaired autophagy at AP circumstance. Within our knowledge, AP-induced dynamic changes of autophagic event influenced by SPINK1 was first observed by the application of this method, though the usage of double-fluorescence construct in monitoring autophagy event of experimental AP was unrare. Other than this, the measurement of neomycin phosphotransferase (GFP-NeoR) fusion protein [12], another substrate of autophagy, was also found to be significantly increased, reconfirmed the presence of hindered autophagy-the turnover blockage of autophagy in the above setting. The observation of impaired autophagy in AR42J-related AP study was consistent to a previous report in which it’s described that the lysosome dysfunction induced impaired autophagy in mouse model of AP [21, 22]. Despite of the aforementioned findings, the pathogenesis of autophagy dysfunction occurred still remains incompletely unclear presently.

Secondly, the relationship between SPINK1 and autophagy was clearly revealed in vitro model of AP. To our knowledge, the inhibitory role of SPINK1 on autophagy in the pancreas was only proved in SPINK1 deficient normal mouse pups[2]. Our investigation, however, just was carried out, possibly for the first time, to set up this connection in vitro study of pancreatic acinar cell of AP. Some of our data partially complied with a previous report in which it states that the inappropriate trypsin activation and pathohistological changes could be partially blocked, and improved in AP setting using SPINK1 transgenic mice[23].

Thirdly, which comes first in the initiation of AP, autophagy or activated trypsin? This cause-effect issue continuously puzzles us and definitely requires an answer. Our data showed that the application of autophagy inhibitors, 3-MA, in our in vitro AP model partially blocked trypsin activity, whereas strengthened autophagy induced by Atg5 transfection upgraded the accumulation of active trypsin in cerulein-manipulated AR42J cells. Therefore, these data suggest that the enhanced autophagy led to trypsin activation in vitro AP. Our data also match in part to a previous report published by Mrs Kugs., et al. in which they claimed that the increased autophagy activity detected in isolated pancreatic acinar cells of cerulein-injected rat upgraded the trypsin activity 48 h after Atg5 transfer [6], though SPINK1’s effects were not studied in his case. Therefore, we concluded that SPINK1 was capable of inhibiting the trypsin activation via amelioration of dysfunctional autophagy in this model. This is significant since SPINK1 has long been considered as a trypsin inhibitor that renders its effects only through their mutual reversible binding, rather than via autophagy inhibition in the pathogenesis of AP. Notably, our data of cellular model need to be interpreted cautiously in the sense that these findings needs to be further approved in in vivo model of AP in the future, though some consistency could be significantly noticed in these data among in vitro and in vivo experiments[7, 23]. However, our results still shed lights on a new way to deeply understand the new mechanism of autophagy through which SPINK1 acts as a trypsin antagonist in acute pancreatitis.

## Acknowledgment

We thank Prof. Chong Yin for providing AR42J cells, Prof. Lin Huang for pMD2.G and psPAX2 packaging plasmids, Associate professor Yang Wang for pCAG-pHluorin-TagRFP-mLC3B plasmid.

